# Aurora A depletion reveals centrosome-independent polarization mechanism in *C.elegans*

**DOI:** 10.1101/388918

**Authors:** K. Klinkert, N. Levernier, P. Gross, C. Gentili, L. von Tobel, M. Pierron, C. Busso, S. Herrman, S. W. Grill, K. Kruse, P. Gönczy

## Abstract

How living systems break symmetry in an organized manner is an important question in biology. In *C. elegans* zygotes, symmetry breaking normally occurs in the vicinity of centrosomes, resulting in anterior-directed cortical flows and establishment of a single posterior PAR-2 domain. Here, we report that zygotes depleted of the Aurora A kinase AIR-1 or of centrosomes establish two posterior domains, one at each pole. Using transgenic animals and microfabricated triangular chambers, we establish that such bipolarity occurs in a PAR-2- and curvature-dependent manner. Furthermore, we develop an integrated physical model of symmetry breaking, establishing that local PAR-dependent weakening of the actin cortex, together with mutual inhibition of anterior and posterior PAR proteins, provides a mechanism for self-organized PAR polarization without functional centrosomes in *C. elegans*.

**One Sentence Summary:** We uncover a novel centrosome-independent mechanism of polarization in *C. elegans* zygotes

---

Symmetry breaking is a fundamental feature of living systems that operates at different scales and contexts. Notably, symmetry must be broken in a tightly coordinated manner at the onset of development to specify embryonic axes. The mechanisms through which this is ensured are incompletely understood.

Symmetry breaking occurs in a stereotyped manner in *C. elegans* zygotes (*1, 2*). Initially, the cell undergoes uniform RHO-1-dependent contractions of the acto-myosin cortex. The RHO-1 guanine-nucleotide-exchange factor (GEF) ECT-2 is then cleared from the cortex close to centrosomes, leading to local cortical relaxation (*3, 4*). Concomitantly, the posterior polarity proteins PAR-2 becomes enriched there, while acto-myosin cortical flows away from this region help segregate PAR-3/PAR-6/PKC-3 towards the future anterior. A partially redundant polarization pathway relies on PAR-2 binding to microtubules, thus shielding PAR-2 from PKC-3-mediated inhibition (*5*). Both pathways are thought to require centrosomes, since laser-mediated centrosome removal early in the cell cycle prevents PAR-2 loading and polarity establishment (*6*). The mechanisms through which centrosomes instruct symmetry breaking remain unclear. Moreover, whereas the system could somehow polarize autonomously has not been assessed.

While investigating the Aurora A kinase AIR-1 during spindle positioning (*7*), we observed that *air-1*(*RNAi*) zygotes exhibited a remarkable polarity phenotype, also noted before (*8*). To investigate this further, we conducted time-lapse microscopy of embryos expressing endogenously tagged NMY-2 to monitor the cortical acto-myosin network, SAS-7 to track centrioles and PAR-2 to assay polarization. By pronuclear meeting, all control embryos established a single PAR-2 domain close to the paternally derived centrioles (Fig. 1A, 1C; MovieS1). In stark contrast, ~80% *air-1*(*RNAi*) embryos formed two PAR-2 domains, one at each pole (Fig. 1B, 1C, MovieS2 “bipolar”); another ~14% established a single PAR-2 domain on the side of the maternal pronucleus (Fig. 1C, “anterior”), with the remaining ~6% exhibiting a single PAR-2 domain on the side of the male pronucleus (Fig. 1C, “posterior”) or no cortical PAR-2 (Fig. 1C, “none”). Analogous distributions were found for endogenous PAR-2 in wild type worms depleted of AIR-1 (Fig. 1D; Fig. S1A), whereas weaker *air-1*(*RNAi*) shifted the phenotype away from bipolar towards one posterior PAR-2 domain (Fig S1B). We conclude that AIR-1, which localizes to maturing centrosomes and also to the cell cortex (*9–11*), ensures uniqueness of symmetry breaking at the presumptive embryo posterior.

**Fig. 1:**
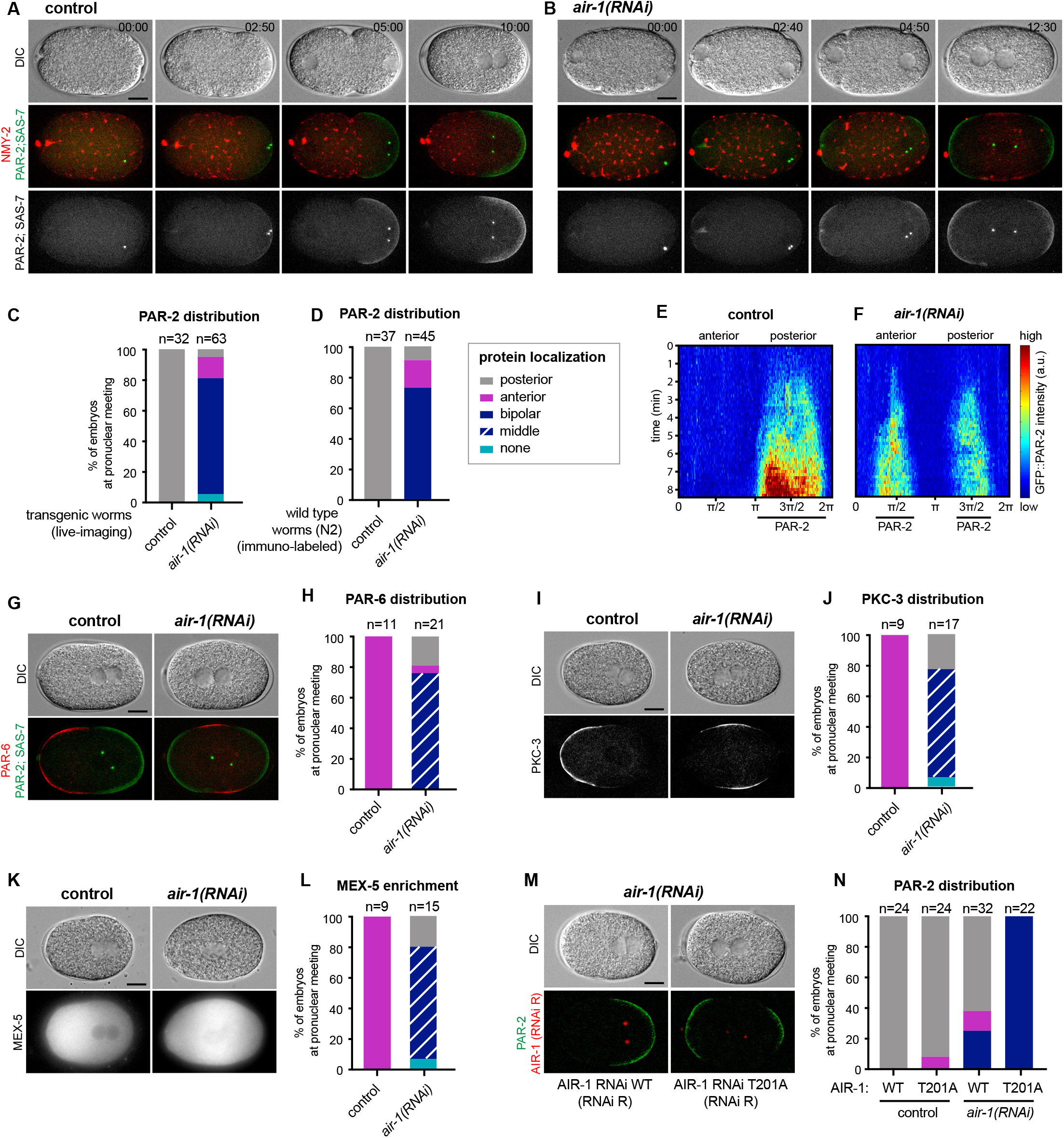
AIR-1 ensures uniqueness of symmetry breaking in *C. elegans* zygotes. (A,B) Time lapse microscopy of control (A) and *air-1(RNAi)* (B) embryos expressing RFP::NMY-2 (red), GFP::PAR-2 and GFP::SAS-7 (both green). In this and all other panels, scale bar is 10μM and time is shown in min:s. (C) Quantification of GFP::PAR-2 distributions at pronuclear meeting. Same timing in all subsequent panels; all statistical analyses (Fischer’s exact test): Table S1. (D) Quantification of endogenous PAR-2 distributions in wild-type (N2) embryos (E,F) Kymograph of GFP::PAR-2 fluorescence along the circumference of control (E, N=12) and *air-1(RNAi)* (F, N=14; bipolar only) embryos. Here and in other kymographs, t=0: onset of cortical flows. (G,J,K) Control and *air-1(RNAi)* embryos expressing mCherry::PAR-2 and RFP::SAS-7 (both green), together with GFP::PAR-6 (red) (G), GFP::PKC-3 (I) or mCherry::MEX-5 (K). (J,L,N) Corresponding quantification of protein distributions. (M) Embryos depleted of endogenous AIR-1 expressing mCh::PAR-2 (red), together with RNAi-resistant GFP::AIR-1 WT (left) or GFP::AIR-1 T201A (right) (both green). (N) Corresponding quantification of GFP::PAR-2 distributions.

We analyzed bipolarization dynamics of *air-1*(*RNAi*) embryos further. The fluorescence intensity of each of the two PAR-2 domains was weaker and their length shorter than that of the single PAR-2 domain in control embryos (Fig. 1E, 1F; Fig. S1C), but their summed fluorescence intensities and lengths were comparable (Fig. S1D, S1E). To assess whether *air-1*(*RNAi*) embryos underwent *bona fide* bipolarization, we examined the distribution of other PAR components. Both PAR-6 and PKC-3, which localize to the anterior in control embryos, were restricted to the middle in ~80% *air-1*(*RNAi*) embryos, as anticipated from PAR-2 bipolarity (Fig. 1G-J). Furthermore, the polarity mediator MEX-5, which is normally enriched in the anterior cytoplasm, was enriched in the middle in ~80% *air-1*(*RNAi*) embryos (Fig. 1K,1L). These findings indicate that the polarization machinery is operational in *air-1*(*RNAi*) embryos, but that its spatial regulation is perturbed.

To determine whether AIR-1 kinase activity is required to ensure uniqueness of symmetry breaking, we used worms expressing RNAi-resistant GFP fusions of either Wild-type or kinase-dead AIR-1(*10*). Wild-type GFP::AIR-1 rescued posterior PAR-2 distribution upon depletion of endogenous AIR-1 in ~63% of embryos, but the kinase-dead version did not (Fig. 1M,N). Therefore, AIR-1 kinase activity is required to ensure unique symmetry breaking. Whereas identification of relevant AIR-1 substrates will be of interest for future investigations, hereafter we utilize this remarkable bipolar phenotype to decipher fundamental principles underlying symmetry breaking at the onset of *C. elegans* development.

Next, we analyzed cortical acto-myosin flows, which normally move away from the centrosome-bearing region (Fig. 2A). By contrast, cortical flows in *air-1*(*RNAi*) embryos stemmed from both poles and were directed towards the center (Fig. 2B). Particle image velocimetry (PIV) of RFP::NMY-2 or of the actin probe GFP::moesin revealed that average peak velocities in control embryos were ~6.8μm/min, but ~3 μm/min on either side upon AIR-1 depletion (Fig. 2B,C; Fig. S2A-C). To address whether this decrease contributes to bipolarity, we co-depleted AIR-1 and the RHO-1 GAP RGA-3, which increases RHO-1 activity and cortical flows (*12*) (Fig. S2, E,F). Although co-depletion augmented flows in *air-1*(*RNAi*) embryos to average peak velocities close to those of control conditions (Fig. 2C), this did not alter the fraction of bipolar embryos (Fig. 2E,F). We conclude that whereas cortical flows are diminished upon AIR-1 depletion, this alone does not explain the bipolar phenotype. To test whether actomyosin flows in *air-1*(*RNAi*) embryos are required for bipolarity, we analyzed *nmy-2(ne3409ts)* embryos, which exhibited severely reduced cortical flows (Fig. S2G). *nmy-2(ne3409)* embryos formed a single posterior PAR-2 domain, albeit smaller and later than in the control (*13*) (Fig. 2G,H). Importantly, depletion of AIR-1 in *nmy-2(ne3409)* embryos almost completely abolished cortical flows; probably as a result, the two PAR-2 domains appeared later and were weaker than in plain *air-1*(*RNAi*) embryos (Fig. 2G, H; Fig. S2G,H,I). These results indicate that some level of cortical flows is needed to establish two robust PAR-2 domains upon AIR-1 depletion.

**Fig. 2:**
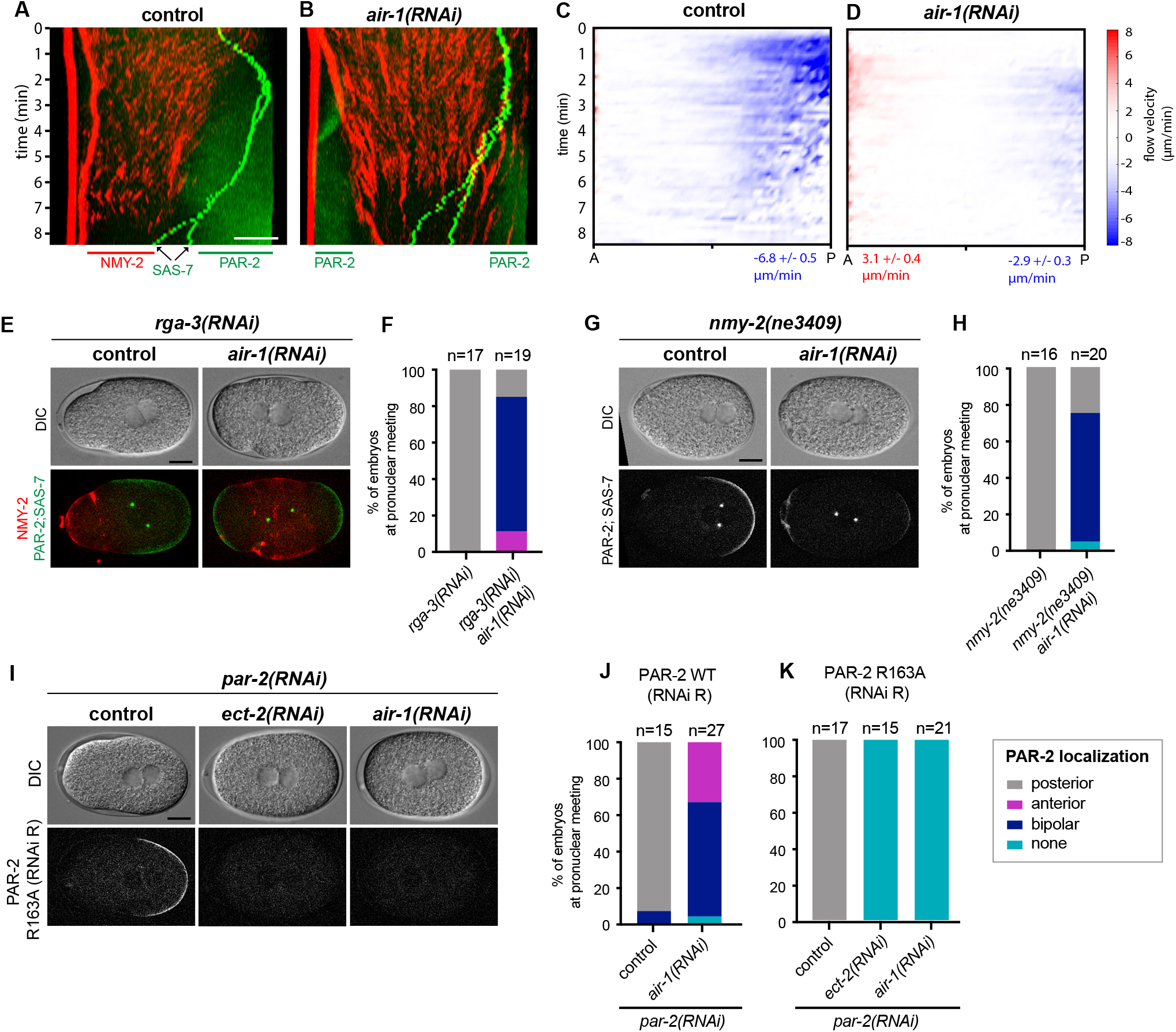
Bipolarity of embryos depleted of AIR-is established through PAR-2. (A,B) Kymographs of control (A) and *air-1(RNAi)* (B) embryos expressing RFP::NMY-2 (red), GFP::PAR-2 and GFP::SAS-7 (both green). Straight red lines: immobile polar bodies. (C,D) Kymograph of cortical flow velocities quantified by particle imaging velocimetry (PIV) in control (C) and *air-1(RNAi)* (D) embryos (N=10 for both). Black lines indicate length of GFP::PAR-2 domain, values the average peak velocities at the anterior (red) and posterior (blue). (E, G) *rga-3(RNAi)* and *rga-3(RNAi)+air-1(RNAi)* embryos (E) or *nmy-2(ne3409)* and *nmy-2(ne3409)+air-1(RNAi)* (G) embryos expressing RFP::NMY-2 (red), GFP::PAR-2 and GFP::SAS-7 (both green). (F, H) Corresponding quantification of GFP::PAR-2 distributions. (I) Embryos expressing GFP::PAR-2 R163A (RNAi-resistant) RNAi-depleted of endogenous PAR-2, together with *ect-2(RNAi)* or *air-1(RNAi).* (J, K) Quantification of GFP::PAR-2 distributions in embryos expressing RNAi-resistant GFP::PAR-2 WT or GFP::PAR-2 R163A, and depleted of endogenous PAR-2 in combination with *air-1(RNAi).*

We set out to test whether bipolarity depends on the partially redundant polarization pathway relying on PAR-2 association with microtubules (*5*). We used a strain expressing an RNAi-resistant GFP tagged version of PAR-2 either wild type (WT) or two mutant forms deficient for microtubule binding (*5*). As reported (*5*), although all three transgenes rescued endogenous PAR-2 depletion, co-depletion of ECT-2 abolished membrane loading of the two PAR-2 mutant forms (Fig. 2I-K; Fig. S2I, K). Strikingly, the same was true when AIR-1 was co-depleted with endogenous PAR-2 (Fig. 2 I-K; Fig. S2J-L). These results demonstrate that both posterior domains in *air-1*(*RNAi*) embryos depend on PAR-2 binding to microtubules.

Since AIR-1 is required for centrosome maturation(*8, 9*), we tested whether depleting other centrosomal components also yielded bipolarity. Strikingly, this was the case in mutants of *spd-2* and *spd-5,* which are required to recruit AIR-1 to centrosomes (Fig. 3A, B), as noted previously(*14, 15*). By contrast, depleting TBG-1, which localizes to centrosomes in an AIR-1-dependent manner (*9*), did not impact polarity (Fig. 3A, 3B) (*9*). Together, these findings indicate that centrosomal AIR-1 is critical to ensure uniqueness of symmetry breaking.

**Fig. 3:**
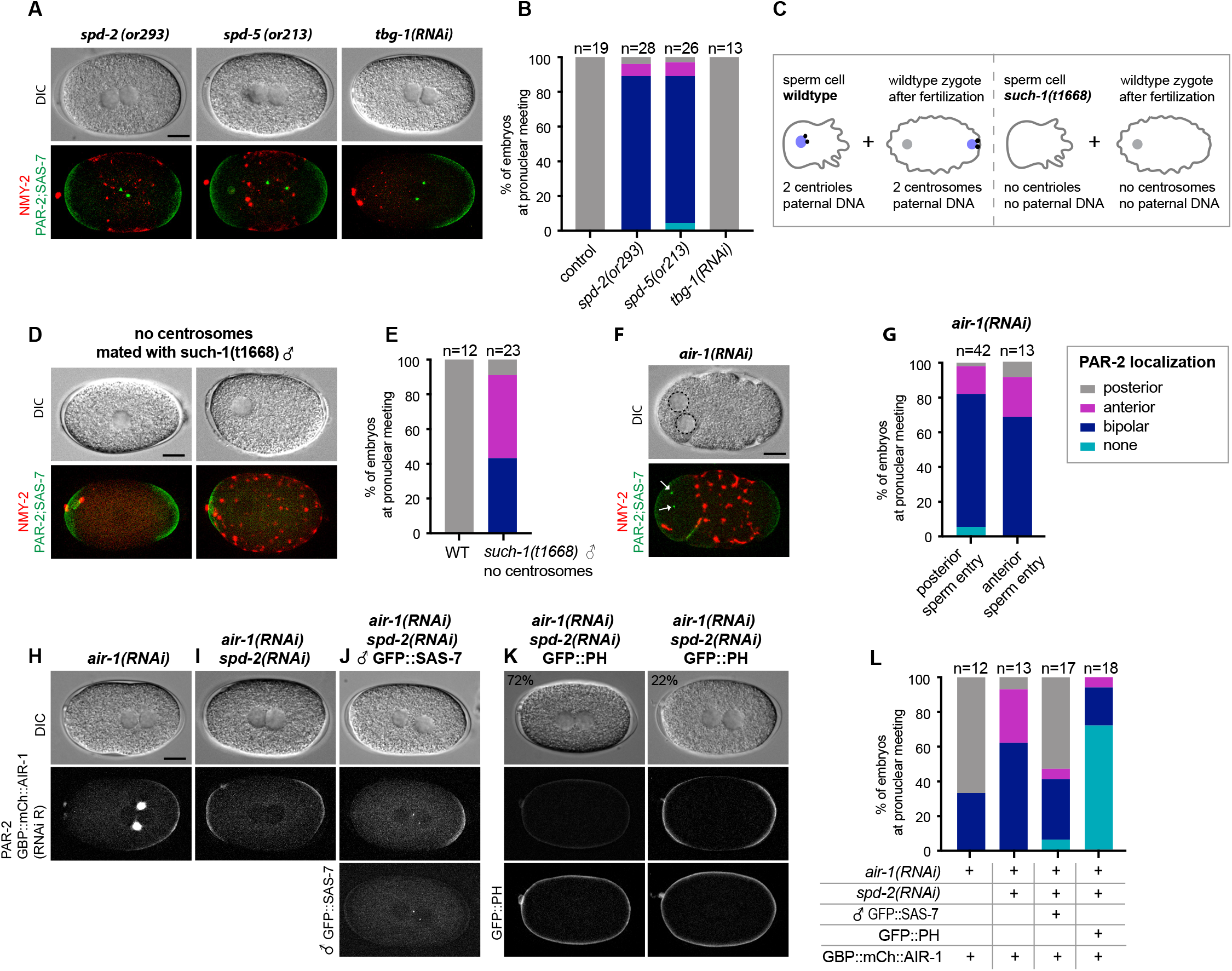
Centrosomes ensure uniqueness of symmetry breaking in *C. elegans* zygotes. (A) *spd-2(or293), spd-5(or213)* or *tbg-1(RNAi)* embryos, as indicated, expressing RFP::NMY-2 (red), GFP::PAR-2 and GFP::SAS-7 (both green). (B) Corresponding quantification of GFP::PAR-2 distributions. (C) Experimental set up for D and E. (D) *fem-1(hcl7)* hermaphrodites expressing RFP::NMY-2 (red), GFP::PAR-2 and GFP::SAS-7 (both green) mated with *such-1(t1668)* males. Note lack of male pronucleus and centriolar GFP::SAS-7. (E) Corresponding quantification of GFP::PAR-2 distributions. (F) *air-1*(*RNAi*) embryo expressing RFP::NMY-2 (red), GFP::PAR-2 and GFP::SAS-7 (both green) with sperm entry next to the maternal pronucleus. Dotted lines: pronuclei; arrows: centrosomes. (G) Quantification of GFP::PAR-2 distributions in *air-1*(*RNAi*) embryos with sperm entry next to maternal pronucleus (“anterior”) or opposite from it (“posterior”). (H-J) Embryos expressing GBP::mCherry::AIR-1 (RNAi-resistant) and mCherry::PAR-2, depleted of endogenous AIR-1 either alone (H) or together with SPD-2 (I,J); in J, hermaphrodites were mated with GFP::SAS-7 males. (K) Examples of embryos expressing GBP::mCherry::AIR-1 (RNAi-resistant), mCherry::PAR-2 and GFP::PH. (L) Corresponding quantification of GFP::PAR-2 distributions.

Considering that centrosomes are thought to be essential for symmetry breaking (*6*), why do *air-1*(*RNAi*) embryos, which do not harbor functional centrosomes, establish two PAR-2 domains? Centrioles in *air-1*(*RNAi*) embryos were usually not coincident with the position of PAR-2 domain initiation (Fig. S3A-C), raising the possibility that bipolarization occurs in a centrosome-independent manner. To test this, we analyzed embryos derived from wild type oocytes fertilized by *such-1(t1668)* mutant sperm, a subset of which does not harbor DNA and centrioles (Fig. 3C) (*16*). Strikingly, embryos fertilized by these sperm, recognizable by the absence of a male pronucleus, microtubule asters and GFP::SAS-7, usually established a bipolar or an anterior PAR-2 domain (Fig. 3D, 3E). Although these experiments indicate that centrosomes are dispensable for symmetry breaking, we considered whether entry of sperm, albeit devoid of DNA and centrioles, might provide an alternative cue. To test this, we took advantage of sperm occasionally entering the oocyte next to the maternal nucleus (*17*). Importantly, we found that most *air-1*(*RNAi*) embryos were bipolar even in this case, indicating that the posterior PAR-2 domain forms independently of the sperm entry site (Fig. 3F, 3G). We conclude that *C. elegans* zygotes undergo unregulated symmetry breaking in the complete absence of centrosomes. The discrepancy with the laser ablation experiments might reflect that lacking centrosomes to start with, as here, or removing them at a later stage, as in the previous work, has distinct consequences.

We sought to test whether centrosomal AIR-1 is sufficient to ensure uniqueness of symmetry breaking. We generated a strain expressing RNAi-resistant AIR-1 fused to mCherry and the GFP binding protein (GBP) to relocalize the kinase. GBP::mCherry::AIR-1 rescued depletion of endogenous AIR-1 in ~67% of embryos, showing that it possesses substantial activity (Fig. 3H, 3L). Co-depletion of endogenous AIR-1 and SPD-2 in this background resulted mainly in bipolar embryos (Fig. 3I, 3L). Strikingly, we found that relocalizing GBP::mCherry::AIR-1 to centrioles using GFP::SAS-7 rescued posterior polarity in most embryos (Fig. 3J, 3L). Importantly, these centrioles do not harbor PCM since SPD-2 is depleted, as evidenced by collapsed spindles (Fig. S3D). We conclude that localizing AIR-1 to centrioles is sufficient to induce a unique posterior symmetry breaking event.

We next utilized GBP::mCherry::AIR-1 to address whether the kinase might play a role also at the cortex, where AIR-1 can be detected (*7*). We examined embryos expressing GBP::mCherry::AIR-1 and GFP::PH, which binds to plasma membrane phosphatidylinositol 4,5-bisphosphate (PI(4,5)P_2_), and depleted of SPD-2 to ensure that no AIR-1 was targeted to centrosomes (Fig. 3K, 3L). Remarkably, forcing GBP::mCherry::AIR-1 to the plasma membrane prevented PAR-2 domain formation in ~72% of embryos, demonstrating that cortical AIR-1 can prevent symmetry breaking. Taken together, these findings lead us to propose that AIR-1 plays a dual function: at the cortex, in preventing erroneous symmetry breaking, perhaps early in the cell cycle, and at centrosomes, in directing symmetry breaking in their vicinity, likely later on.

Why do embryos become bipolar upon AIR-1 depletion or without centrosomes? Because PAR-2 domains developed almost invariably at the poles, we hypothesized that membrane curvature might be important. To test this, we microfabricated equilateral triangular PDMS chambers ~40 μm in side length and ~ 25 μm in depth, in which embryos were squeezed. Despite being deformed, control embryos established a single PAR-2 domain in the vicinity of centrioles, which spread along one side or in one corner (Fig. 4A-D, MovieS3). In contrast, in *air-1*(*RNAi*) embryos, one or sometimes two PAR-2 domains formed almost invariably in one of the corners, independently of the position of centrioles or polar bodies marking the position of the maternal pronucleus (Fig. 4A-D, MovieS4,5). Therefore, without functional centrosomes, PAR-2 domains form preferentially in regions of high membrane curvature.

**Fig. 4:**
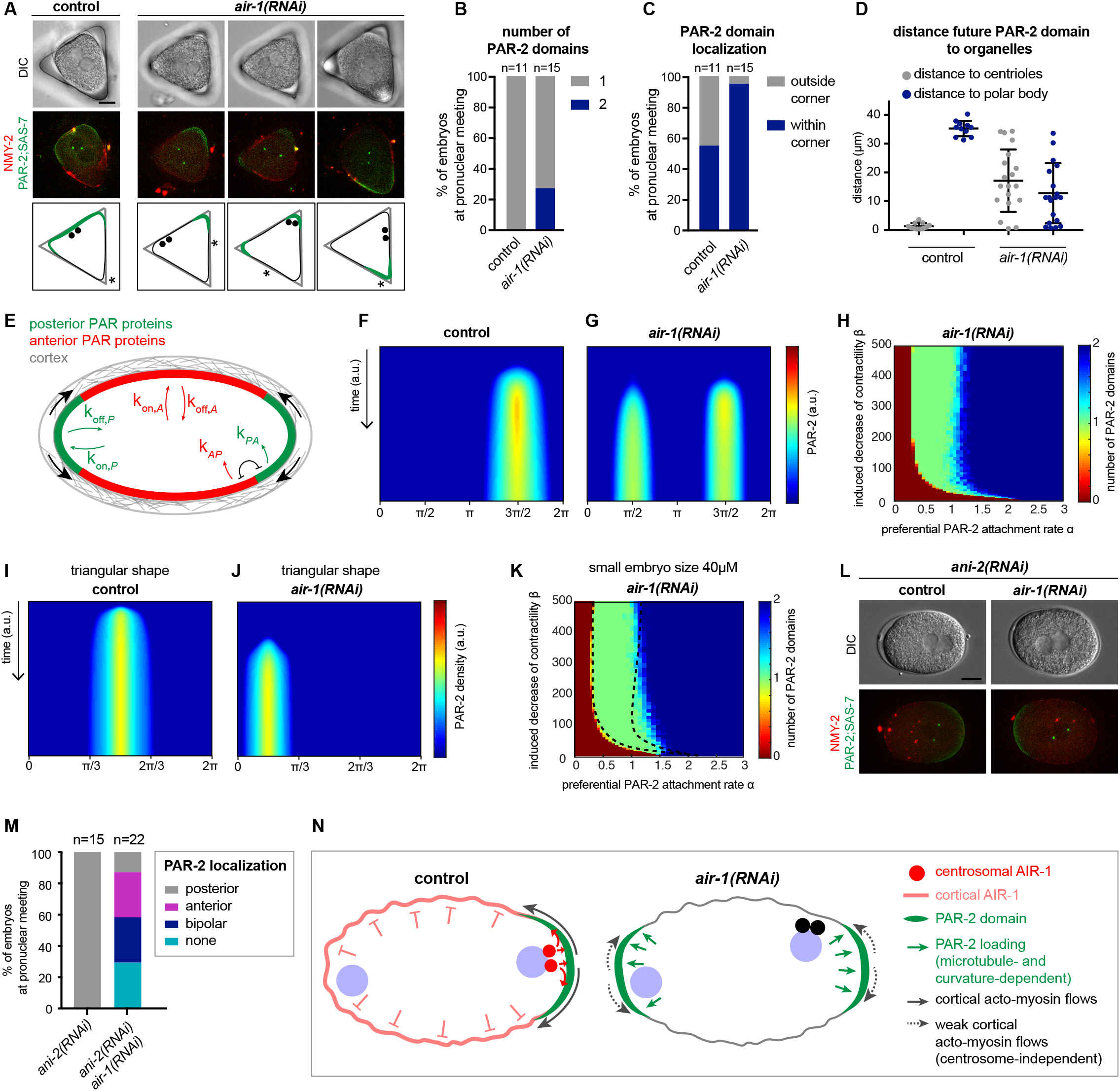
Physical model of symmetry breaking without centrosomes in areas of high membrane curvature. (A) Control and *air-1*(*RNAi*) embryos expressing RFP::NMY-2 (red), GFP::PAR-2 and GFP::SAS-7 (both green) placed into triangular PDMS chambers 40μM in side length. Lower panels illustrate the localization of PAR-2 domains (green), centrioles (black dots) and polar bodies (black arrow). (B-D) Corresponding quantification of GFP::PAR-2 distributions (domain number, B; domain localization, C; domain distance to centrioles and polar bodies, D). (E) Principal tenets of physical model; see Supplementary Information for details. (F,G) Simulation kymographs of PAR-2 density in control (G) and *air-1(RNAi)* (H) embryos. (H) Phase diagram showing co-existence of polarity states in *air-1(RNAi)* embryos (regular size: 57 μM). (I,J) Simulation kymographs of PAR-2 distribution in control (J) and *air-1(RNAi)* embryos (K) in triangular chambers. (K) Phase diagram showing co-existence of polarity states in *air-1(RNAi)* for 40μM-long embryos; dotted lines indicate bipolar:monopolar boundary in 57μM-long embryos (see H). (L) *ani-2(RNAi)* and *ani-2(RNAi) +air-1(RNAi)* embryos expressing RFP::NMY-2 (red), GFP::PAR-2, GFP::SAS-7 (both green). (M) Corresponding quantification of GFP::PAR-2 distribution. (N) Working model of centrosome-independent symmetry breaking in *C. elegans.* See text for details.

To further analyze how membrane curvature cue can lead to bipolarization, we developed a physical model (Fig. 4E, Supp. Mat.). This model is based on known interactions between anterior and posterior PAR proteins (*18*) and accounts for preferential PAR-2 binding to regions of high curvature, as well as for coupling between PAR-2 and cortical actin. In this model, the wild type situation is simulated by locally weakening acto-myosin activity next to centrosomes (Fig. 4F; Fig S4A, MovieS6). Independently of the location of the weakening, a posterior domain always establishes at one pole. Importantly, if acto-myosin activity is not weakened locally, as without functional centrosomes or upon AIR-1 depletion, then the corresponding phase diagram exhibits regions of coexisting polar and bipolar states (Fig 4G, 4H; Fig S4B, MovieS7). Moreover, the fact that one domain forms more readily than two domains in the triangular geometry is captured by the model (Fig.4I, 4J and Fig. 4A-D). We explored the validity of the physical model further by testing the impact of smaller embryo size, for which the model predicts a shift towards a single PAR-2 domain in *air-1*(*RNAi*) embryos (Fig. 4K). Accordingly, we found that smaller *ani-2(RNAi)* embryos (*19*) are bipolar in only ~29% of cases, with ~ 43% harboring a single PAR-2 domain, with a bias towards the anterior side (Fig. 4L,4M; Fig. S4C). Perhaps this bias arises from the maternal meiotic spindle being anteriorly localized, since this can promote PAR-2 loading in some circumstances (*20*). We conclude that our integrated physical model accounts for polarization in *C. elegans* zygotes in the absence of functional centrosomes, and supports the notion that the system can self-organize polarity in an autonomous manner.

In conclusion, our work reveals the mechanism through which AIR-1 normally orchestrates polarity establishment in *C. elegans* zygotes, and uncovers self-organizing properties that polarize embryos when centrosomes are missing, which could be of particular importance to establish embryonic axes in parthenogenetic species lacking centrioles. We propose that *C. elegans* AIR-1 normally acts through a dual-mechanism: prevention of erroneous symmetry breaking events through cortical localization and induction of a single symmetry breaking via centrosomal localization (Fig. 4N). In doing so, AIR-1 funnels the inherent self-organizing properties onto a single site, next to paternally contributed centrioles, thus ensuring spatial coupling between paternal and maternal contributions at the onset of development. It will be interesting to investigate whether such a dual-function extends to the human homologue Aurora A oncogene in polarized tumorigenic settings (*21*).

## Acknowledgements

We are grateful to Carrie Cowan and Sachin Kotak for sharing unpublished observations and fruitful discussions, as well as to Bruce Bowerman, Fumio Motegi, Asako Sugimoto and Geraldine Seydoux for their gift of worm strains, as well as the *Caenorhabditis* Genetics Center (CGC), which is funded by NIH Office of Research Infrastructure Programs (P40 OD010440). We thank **Alexandra Bezler, Alessandro De Simone and Radek Jankele for comments on the manuscript.** We thank the Microstructure Facility (CMCB, TU-Dresden), in part funded by the State of Saxony and the European Fund for Regional Development — EFRE, for the production of the micro-well structure. This work was supported by post-doctoral fellowships from EMBO to Ke.Kl. (ALTF 81-2017) and to M.P. (ALTF 1426-2016), the Fondation Bettencourt.Schueller prize to N.L., as well as grants from the Swiss National Science Foundation to Pi.Gö. (31003A_155942) and to Ka.Kr., (205321_175996), and from the European Research Council (281903 and 742712) to S.W.G. and Pe.Gr.

## Author contributions

Ke.Kl. and Pi.Gö. designed the project; Ke.Kl. conducted experiments with support from L.v.T., S.H. and C.B.; Ke.Kl. and Pi.Gö. analyzed the data; N.L. and Ka.Kr. developed the physical model; S.W.G. and Pe.Gr. developed the PDMS chambers; M.P. and C.G. generated worm strains; Ke.Kl., Pi.Gö., N.L. and Ka.Kr wrote the manuscript. We have no competing interests to declare.

## Data and materials availability

All data is available in the manuscript or the supplementary materials.

## Supplementary Materials

**Material and Methods**

**Supplementary References**

**Supplementary Figures S1-S4**

**Supplementary Tables:**

Table S1: list of worm strains

Table S2: statistical analysis of presented bar graphs

## Supplementary Movies

Movie S1: control

Movie S2: *air-1(RNAi)*

Movie S3: triangle control

Movie S4: triangle *air-1(RNAi)*

Movie S5: triangle *air-1(RNAi)* _2

Movie S6: model control

Movie S7: model *air-1(RNAi)*

**Movie S1,S2:** Embryo expressing RFP::NMY-2 (red); GFP::PAR-2 and GFP::SAS-7 (both green). Left: DIC, right: merge. Control (Movie S1), *air-1(RNAi)* (Movie S2). In all movies, elapsed time is shown in min:s.

**Movie S3-S5:** Embryo placed in a triangular chamber expressing RFP::NMY-2 (red); GFP::PAR-2 and GFP::SAS-7 (both green). Left: DIC, right: merge. Control (Movie S3), *air-1(RNAi)* (Movie S4,S5).

**Movie S6,S7:** Simulation of density of anterior and posterior PAR proteins (aPAR, pPAR) over time using physical model. Control (Movie S6), *air-1(RNAi)* (Movie S7).

## Materials and Methods

### Worm Strains

Nematodes were maintained at 24°C using standard protocols (Brenner,1974). Worms carrying temperature-sensitive mutations were maintained at permissive temperature (16°C) and shifted to restrictive temperature (24°C) for 24h prior to imaging. See Table S1 for the list of worm strains used in this study. New transgenic worm strains were generated as follows:

The GBP::mCherry::AIR-1 construct was obtained by inserting the mCherry coding sequence at the 3’-end of the GFP-binding protein (GBP), obtained from the pAOD-VHHGFP4 vector (a kind gift from Aurelien Olichon), followed by codon optimized RNAi-resistant *air-1* cDNA (a kind gift from Asako Sugimoto). Flexible linkers of thirteen amino acids (GAGAGAGAGAFSV) were inserted between GBP and mCherry as well as between mCherry and air-1 cDNA. The corresponding transgenic line was generated by microparticle bombardment of *unc-119(ed3)* mutant worms (*22*).

CRISPR/Cas9-mediated genome editing was used as described previously (*23*) to insert tagRFP-T in frame at the 5’ of *sas-7* locus. To target Cas9 to the *sas-7* locus, the 5’gcttaaaatcaactcaccg(TGG)3’ −19bp targeting sequence was inserted into the Cas9-sgRNA construct (pDD162). Homologous repair template for insertion of the self-excising selection cassette (SEC) was generated by modifying pDD284 vector using Gibson assembly. Left and right homology arms of 655bp and 652bp respectively were PCR-amplified and inserted into pDD284 opened using Clal, Spel and Psil to remove flag tag from the original vector. A silent mutation in the PAM motif (5’TGG3’ --> 5’TAG’) was introduced to prevent repair template cutting and a 5’CACTCCACTGGAACCTCTAGA3’-21bp linker was added in between *tagRFP* and *sas-7* coding sequences. An injection mix containing 50ng/μl targeting vector, 50ng/μl homologous repair template, 10ng/μl pGH8 (*Prab-3::mcherry::unc-54 3’UTR*), 5ng/μl pCFJ104 (*Pmyo-3::mcherry::unc-54 3’UTR*) and 2.5ng/μl pCFJ90 (*Pmyo-2::mcherry::unc-54 3’UTR*) was micro-injected into the gonads of N2 worms. Selection of insertion events and removal of the SEC cassette was performed as described (*23*).

A similar strategy was used to generate endogenously RFP-tagged SPD-2 expressing line. The 5’tgttcattacagagattcat(TGG)3’-21bp targeting sequence was cloned into pDD162 vector and pDD286 was modified using Gibson assembly and Q5-directed mutagenesis to generate the homologous repair template. PCR amplified 639bp-left and 667bp-right homology arms were inserted into pDD286 digested with ClaI and NgoMIV and 6 silent mutations were introduced in the targeted sequence and the PAM site 5’tg<u>c</u>tc<u>g</u>tt<u>g</u>ca<u>c</u>ag<u>g</u>ttcat(<u>G</u>GG)’3. As before, an appropriate injection mix was micro-injected in N2 worm’s gonads and screening was performed as described (*23*).

### RNAi

RNAi-mediated depletion was performed essentially as described (*24*), using bacterial feeding strains from the Ahringer (*25*) or the Vidal library (*26*) (the latter a gift from Jean-François Rual and Marc Vidal). The bacterial feeding strain for depleting specifically endogenous AIR-1 without affecting *air-1* RNAi resistant transgenes (*tjlsl73* and *tjlsl88*) expression was a kind from Asako Sugimoto (*9*). The bacterial feeding strains used to deplete specifically endogenous PAR-2 in strains expressing RNAi-resistant par-2 transgenes (*axIs1933, axIs1936, axIs1934*) was a kind gift from Fumio Motegi and Geraldine Seydoux (*5*). RNAi for *air-1* (Vidal), *rga-3* (Vidal), *spd-2* (Vidal), *ani-2* (Ahringer), *c27d9.1* (Vidal) was performed by feeding L3-L4 animals with bacteria expressing the corresponding dsRNA at 24°C for 16-26 hours. For all experiments with double RNAi, single RNAi was diluted 1:1 with bacteria expressing an empty vector (L4440). In Fig. S1B, *air-1(RNAi)* was diluted 1:5 or 1:10 with bacteria expressing an empty vector (L4440). RNAi for *tbg-1* (Ahringer) and *tbg-1+air-1* was performed by feeding L2-L3 animals with bacteria expressing dsRNA at 20°C for 48 hours and imaging embryos of their offspring. Strains carrying temperature sensitive mutations were shifted to the restrictive temperature (24°C) for 24h and RNAi performed as described above.

### Live imaging

Gravid hermaphrodites were dissected in osmotically balanced blastomere culture medium (Shelton and Bowerman, 1996) and the extracted embryos mounted on a 2% agarose pad. Fluorescence and DIC time lapse microscopy was performed at room temperature with a 60x CFI Plan Apochromat Lambda (NA 1.4) objective on Nikon Eclipse Ti-U Inverted Microscope connected to a Andor Zyla 4.2 sCMOS camera or with a 63x Plan-Apochromat (NA 1.4) objective on a Zeiss ObserverD.1 inverted microscope connected to the same type of camera. Embryos were imaged with a frame rate of 1 frame every 10 seconds and a z-stack was acquired at every time point covering 20μm, with a distance of 0.7μm between focal planes. Timelapse images in Fig. S2A,B were acquired at 23°C using an inverted Olympus IX 81 microscope equipped with a Yokogawa spinning disk CSU - W1 with a 63x (NA 1.42 U PLAN S APO) objective and a 16-bit PCO Edge sCMOS camera. Images were obtained using a 488 nm solid-state laser with an exposure time of 400 ms and a laser power of 60%. Embryos were imaged with a frame rate of 1 image every 5 seconds a z-stack was acquired at every time point covering 2μm of the embryo cortex with a z-step size of 0.5 μm between focal places.

### Image processing and analysis

Images acquired at the spinning disk were processed by z-projecting the 4 cortical planes using maximum intensity (ImageJ software). Images acquired with the epifluoresence microscope (Zeiss ObserverD.1or Nikon Eclipse Ti-U) were processed as follows using ImageJ software: images underwent background subtraction using a “rolling ball” algorithm of 10 pixels, followed by a maximum intensity z-projection. Grey levels were set identically for all images within each experimental series.

For flow velocity analysis, heat maps were obtained using Particle Image Velocimetry (PIV) with a freely available PIVlab MATLAB algorithm (pivlab.blogspot.de). Using PIVlab, we performed a 4-step multi pass with a final interrogation area of 8 pixels with a step of 4 pixels. 2D velocity fields were obtained by averaging the x-component of velocity along the y-axis for each value in a single frame. All values were averaged over several embryos and represented in a heatmap using MATLAB. Prior to averaging all values across multiple embryos, embryos were aligned temporally using the best fit of the pronuclei growth curves. Pronuclear diameter was measured manually using ImageJ on DIC images.

For Fig. S3C centrosomes were tracked in 3 dimensions using the freely available ImageJ plugin TrackMate (imagej.net/TrackMate).

### PAR-2 domain extent and intensity measurements

Images were processed as described above using ImageJ, following straightening of the embryo’s outline for each time point using the ImageJ plugin “Straighten” (imagej.nih.gov/ij/plugins/straighten.html). The intensity of each pixel along the embryo’s straightened circumference was measured with ImageJ, and the resulting values displayed in a heatmap representing the GFP::PAR-2 intensity using Matlab.

To calculate the extent of the PAR-2 domain, the circumference of the cell was smoothened for each time point using a 23-point moving average to reduce acquisition noise in Matlab. For each time point, a noise floor estimation was obtained by detecting the max and min values following a smoothing by applying a moving average. The noise floor value was then approximated using the minimum values. The value of the average background noise was used as threshold value to determine the points to constitute the domain boundary. The same threshold was applied to all analyzed image sequences. In order to reduce noise-induced threshold crossing, the data values were squared.

### Mating experiments

SUCH-1 is a component of the anaphase promoting complex/cyclosome (APC/C) and its depletion causes, in addition to the paternal sperm phenotype, a maternally contributed delay in mitosis. Therefore, we used *such-1(t1168)* males to fertilize wild type females *(fem-1(hc17)),* thus ensuring that all embryos resulted from *such-1(t1168)* mutant sperm and that the maternal contribution is normal. To this end, gravid *fem-1(hc-17)* hermaphrodites expressing RFP::NMY-2; GFP::PAR-2; GFP::SAS-7 were shifted to the restrictive temperature (24°C) and L4 progeny were then mated with *such-1(t1668)* males at 20°C for 24h. Gravid adults were dissected and embryos were imaged as described above. Prior to and after imaging, each embryo was thoroughly screened for the absence of centrosomes as evaluated by the absence of focal GFP::SAS-7 signal.

To target GBP::mCherry::AIR-1 to centrosomes, L4 larvae expressing mCherry::PAR-2 and GBP::mCherry::AIR-1 were mated with GFP::SAS-7 males for 24h at 20°C on *spd-2(RNAi).* Gravid adults were dissected and embryos imaged as described above. Only embryos that were fertilized, as evaluated by the presence of GFP::SAS-7, were imaged.

### Immunofluorescence

Embryos were permeabilized by freeze-cracking followed by fixation in methanol at −20°C for 5 min, followed by incubation with primary antibodies (mouse anti-α-tubulin (1/200; DM1A, Sigma) and rabbit anti-PAR-2 (1/200) (*27*) for 1 h at room temperature. Secondary antibodies were Alexa-Fluor-488-coupled anti-mouse-IgG and Alexa-Fluor-568-coupled anti-rabbit-IgG, both used at 1:500. Slides were counterstained with 1 mg/ml Hoechst 33258 (Sigma) to reveal DNA.

### Triangular PDMS chambers

Micro-well structures were produced using standard soft-lithography methods. Using a custom lithography mask, a silicon waver coated with AZ 15nXT (MicroChemicals GmbH, Ulm, Germany) was illuminated with UV light to generate a casting mold of triangles with a side length of 40μm. Micro-well structures were generated by pouring poly(dimethylsiloxane) (PDMS; Sylgard 184, VWR) into these casting molds. Finally, these PDMS structures were cured for 1.5 hours at 75°C. Before each experiment PDMS chambers were treated with oxygen plasma for 90s to reduce its hydrophobicity. Worms were dissected as described above, and embryos were placed into the triangular chamber with an eyelash tool prior to symmetry breaking. Epifluoresence imaging was performed as described above.

### Statistical analysis

Statistical significance for all shown quantification was performed using the non parametric two-sided Fisher’s exact test, comparing a single category against the other pooled categories (e.g. posterior vs non-posterior) for two conditions (e.g. control versus air-1(RNAi)). The corresponding p-values for all experiments can be found in Table S2. A p-value <0.05 was considered as statistically significant.

### Theoretical description of symmetry breaking with and without external cue

#### A) Description of the framework

The process of polarization depends on competitive binding of anterior and posterior proteins to the plasma membrane coupled to cortical actin flows. These processes can be described by a non-linear advection-reaction-diffusion system (*18*). This has been found to successfully describe polarization of wild-type embryos when an appropriate cortical actin flow is imposed, reflecting the centrosome-derived cue. The dynamic equations in this system for the concentrations *A* and *P* of membrane-bound anterior (PAR-6) and posterior (PAR-2) proteins, respectively, are

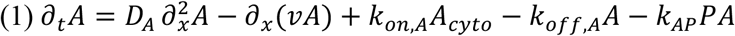

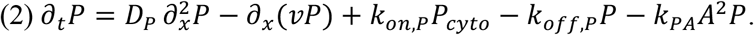

Here, *D_AP_* denote the diffusion coefficients of the two species, *ν* the velocity of the cortical flow, *k_on_* and *k_off_* the respective attachment and detachment rates, and *A_cyto_* and *P_cyto_* their cytoplasmic concentrations. The latter are equal to the total number of proteins minus the membrane bound proteins divided by the embryo volume. The last terms *k_AP_PA* and *^k^PA*^*A*^2^*P*^ account for the mutual antagonism between anterior and posterior PAR proteins (*18*). In the equations, we exploit that the embryo is symmetric with respect to rotations around the long axis and describe the concentrations only as a function of the remaining coordinate along the membrane, which is denoted by *x*. We scale lengths, such that 0 ≤ *x* ≤ 2*π*. The equations are complemented by periodic boundary conditions. Then, cytoplasmic and membrane concentrations of the anterior proteins are related through 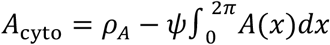 where *ρ_A_* is the total amount of proteins divided by the volume of the embryo (i.e. cytoplasmic concentration if there is no protein on the membrane), and *ψ* the area/volume ratio of the embryo.

The reaction kinetics can exhibit bi-stability, such that for a specific set of parameter values either an *A*-dominant state with high *A* and low *B* or a *P*-dominant state with high *P* and low *A* is reached. There is also an unstable state with intermediate concentration of both *A* and *P*. These homogenous, unpolarized states give way to a polarized state if a strong enough flow is applied, with one part of the embryo being in the *A*-dominant state (anterior domain) and the other in the *P* -dominant state (posterior domain). Once the polarized state is established, it persists even when the flow ceases (*18*).

An experimentally constrained set of parameters for which the above behavior holds has been determined experimentally (*18*). Here, we will use similar values (see Table in C, below) with one important modification: our experimental observations suggest that the attachment rate of the posterior proteins is higher in regions with larger curvature. We account for this observation by imposing 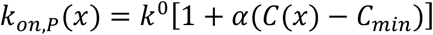, where *C* is the membrane curvature at position *x*. Furthermore, *k*^0^ is the rate of attachment to the minimally curved part of the embryo and *a* quantifies the coupling between the “excess”-curvature and the PAR-2 attachment rate. For *k*^0^ we take the value obtained by Goehring et al. (*18*).

The velocity *ν*, of the cortical actin flow remains to be specified in this novel framework. In most previous work, flows were assumed to be directly linked to the action of centrosomes at the posterior pole and imposed *ad hoc*, even though the exact nature of this action and its temporal regulation remain unclear (*18, 28, 29*). In AIR-1 depleted embryos, weak cortical flows emanating from both poles can be observed despite non functional centrosomes, and it remains unknown how these cortical flow are initiated. We complement the above equations with an explicit description of the actin dynamics, which will notably yield an actin flow. Similar to Ref. (*30*), we base our description on actin conservation and force balance. Explicitly,

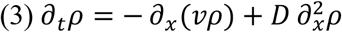

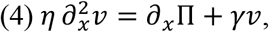

where *ρ* is the cortical acto-myosin gel density, *D* an effective diffusion constant,*η* its viscosity, and *γ* quantifies the gel friction against the membrane. The function Π denotes an effective cortical stress and, in particular, accounts for motor-induced contractility. Gradients in Π along the membrane generate flows over a typical distance 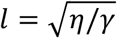. Following (Joanny et al.) (*30*), we choose Π = −*aρ*^3^ + *bρ*^4^ with *a* and *b* > 0 so that the cortex is contractile for moderate densities and that the common positive pressure is recovered for high densities. The experiments suggest that PAR-2 loading affects actin mechanics; in particular, there are no flows in the absence of PAR-2 and AIR-1 (Fig. S2N). We assume that cortex contractility decreases with increasing PAR-2 concentration and account for this by setting *a* = *a*^0^ − *βP*. In contrast, we choose the passive coefficient *b* to be uniform. In our phenomenological description, we do not specify a molecular mechanism through which *P* reduces contractility, as this is irrelevant for the qualitative behavior of the system. Previous work suggests that posterior PAR proteins decrease the concentration of active myosin through the RHO-1 pathway. However, since we focus here on the essential parts of the physical mechanism underlying symmetry breaking, we chose not to include these molecular details. Likewise, we do not consider the actin network and the myosin distributions separately, even though this distinction might be relevant in other situations.

Similar to equations (1) and (2), the full set of equations (1)-(4) has an *A*-dominant and a *P*-dominant stationary state. Due to the dependence of *P* binding on the curvature, however, they are not completely homogenous. For a sufficiently strong preferential attachment rate *α* or decrease of contractility *β*, these states are not stable: *P* will bind preferentially to the membrane in a curved region, which generates a small gradient of *P* along the membrane. This protein gradient leads to a gradient of contractility, which in turn generates an actin flow directed away from

The wild-type situation is obtained by locally weakening the cortex at the posterior pole, mimicking the centrosomal cue, which induces cortical flow from posterior to anterior pole. This is achieved by initially disrupting the cortex (that is setting *ρ* = 0) at the posterior pole. We get qualitatively similar results by reducing activity (e.g. setting *a* = 0) at this pole. Then, embryos always develop a posterior PAR-2 domain, similar to what is described in (Goehring et al) (*18*) (where the flow is inserted by hand). Strikingly, our model constrains the PAR domains to lie at poles of the embryo, even if the cue is not delivered on a pole. Moreover, if we shift all densities of the polarized state by at most ±*π*/2 along *x*, the system spontaneously returns to its original state. This result, mimicking the *in-vivo* experiment of Mittasch et al. (*31*), offers a way to understand the robust positioning of the PAR proteins at the cell poles.

##### Typical *air-1*(*RNAi*) embryo

We first consider a typical embryo of length 57 μm (*18*) and aspect ratio 1.6 without centrosomal cue. Results of varying *α* and β are given in Fig. 4I. For small *α* or small *β*, the embryo never polarizes and stays in an *A*-dominant state. Increasing a for a given *β*, we get first polar embryos (anterior or posterior). Increasing a further leads to coexistence of phases where either polar or bipolar embryos are obtained, depending on the initial random condition. Finally, increasing a further still leads to 100% of bipolar embryos. Kymographs (Fig. 4G,H) were obtained with *α* = 1.2 and β = 100.

##### Smaller embryo

When considering an embryo with the same aspect ratio and a reduced length of 40 μm, one obtains the phase diagram shown in Fig. 4L. While qualitatively similar to the one above, it is clear that the polar phase (green) is larger, with the frontier with the bipolar phase shifted to the right. Hence, some embryos that were mainly in the bipolar phase are now in the coexistence of the polar and bipolar phases, in agreement with the experimental results. This shift is due to two effects. First, the curvature difference between the poles and the equator is higher, so that the enhanced rate of PAR-2 attachment at poles is higher. This effect globally shifts phases to the left. Note that we get a similar effect by increasing the aspect ratio. The other dominant effect is due to the reduced area/volume ratio, which enables less PAR-proteins to attach to the membrane, and consequently diminishes the coupling with acto-myosin contractility. This globally moves phases to the top of the phase diagram.

##### Triangles

Finally, we apply our description to embryos squeezed in triangular chambers. Contour length is now approximately 120 μm, and the area/volume ratio is 0.22*μm*^-1^. We model the curvature function as *C*(*x*) = *C*_0_[1 − 0.6cos(3*πx/L*)^2^]^-6^ with *C*_0_ = 0.05*μm*^-1^. This gives rise to phenotypes with one, two or three domains, depending on the parameters a and *β*. Due to increased PAR-2 domain extension in this triangular shape the bipolar phenotype appears to be unstable, leading to a shift towards a single PAR-2 domain in this scenario. Kymographs in Fig.4 J,K were obtained with *α* = 0.9 and *β* = 20.

#### C) Parameters

Parameters for the advection-reaction-diffusion system are chosen following Goehring et al. (*19*). The (unknown) rates of mutual antagonism *k_AP_* and *k_PA_* are chosen such that the system is in the bi-stable phase where !-dominant and polarised states coexist and are stable (phase (ii) (*19*)).

The hydrodynamic length of the actin cortex is set to 15 μm, following (Mayer et al.) (*32*). The effective diffusion constant D is obtained by studying experimental results for the steady-state actin density in a WT-polarized state, balancing advective and diffusing flux. We get *D* ≃ 0.3*μm*2/*s*. Finally, parameters *a* and *b* remain unknown but do not affect qualitatively the resulting phenotypes. We chose a set such that a uniform actin cortex is stable, and that the typical velocity magnitude of cortical flows is 4 μm/min.

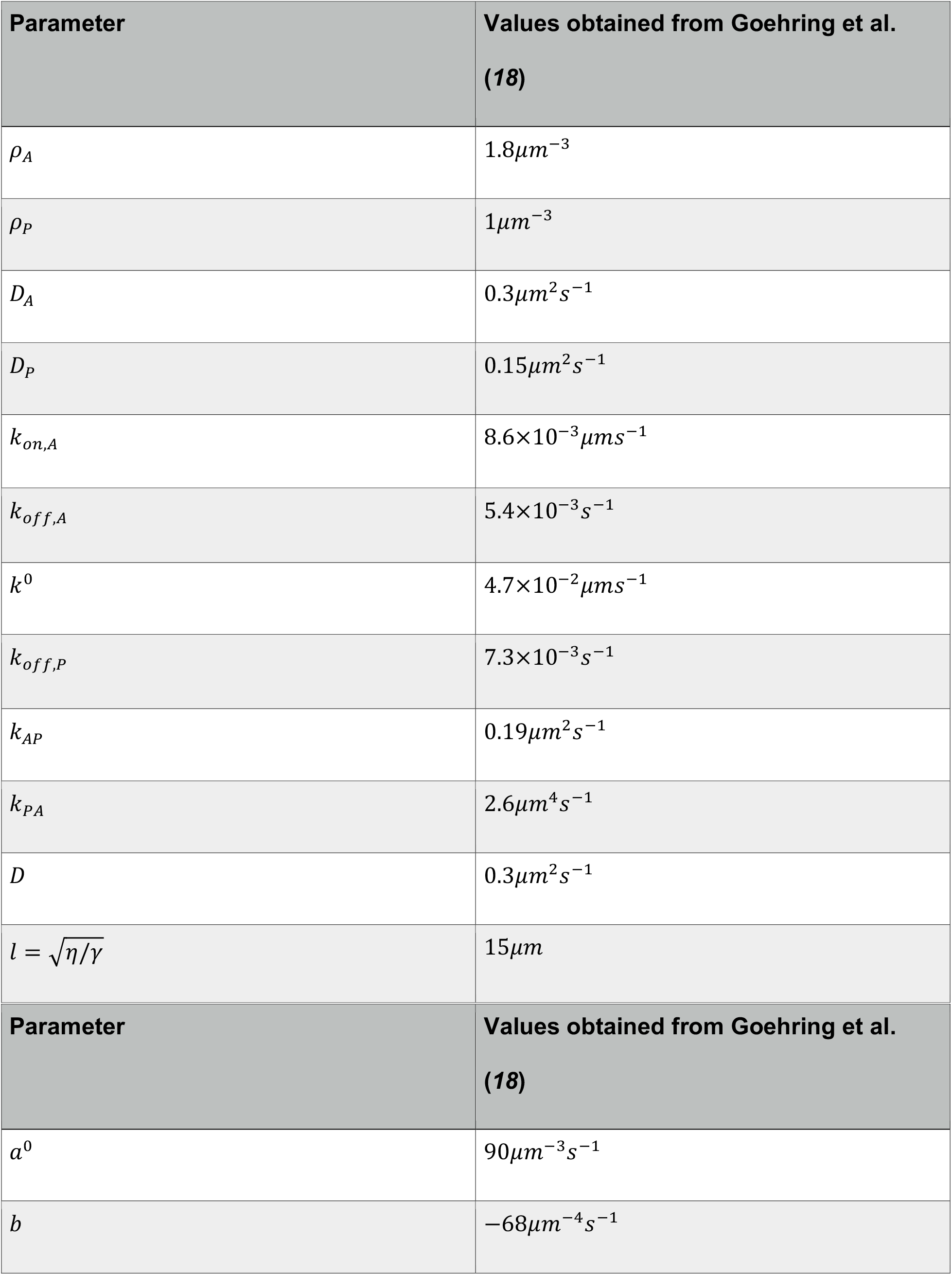

| Parameter | Values obtained from Goehring et al.<br>(18) |
| --- | --- |
| $\rho_A$ | $1.8\mu\text{m}^{-3}$ |
| $\rho_P$ | $1\mu\text{m}^{-3}$ |
| $D_A$ | $0.3\mu\text{m}^2\text{s}^{-1}$ |
| $D_P$ | $0.15\mu\text{m}^2\text{s}^{-1}$ |
| $k_{on,A}$ | $8.6\times 10^{-3}\mu\text{m}\text{s}^{-1}$ |
| $k_{off,A}$ | $5.4\times 10^{-3}\text{s}^{-1}$ |
| $k^0$ | $4.7\times 10^{-2}\mu\text{m}\text{s}^{-1}$ |
| $k_{off,P}$ | $7.3\times 10^{-3}\text{s}^{-1}$ |
| $k_{AP}$ | $0.19\mu\text{m}^2\text{s}^{-1}$ |
| $k_{PA}$ | $2.6\mu\text{m}^4\text{s}^{-1}$ |
| $D$ | $0.3\mu\text{m}^2\text{s}^{-1}$ |
| $l = \sqrt{\eta/\gamma}$ | $15\mu\text{m}$ |
| $a^0$ | $90\mu m^{-3}s^{-1}$ |
| $b$ | $-68\mu m^{-4}s^{-1}$ |

#### D) Simulations

Equations (1)-(4) are solved numerically using a finite-difference scheme. In each time-step, cortical velocity is obtained in Fourier space by solving (4) and back-transformed to real space. Then, actin density and protein densities are updated in Fourier space using Fourier-transform of (*1–3*), and then back-transformed to real space. The spatial discretization step is 0.7 μm.

## Supplementary Figures

**Fig. S1:**
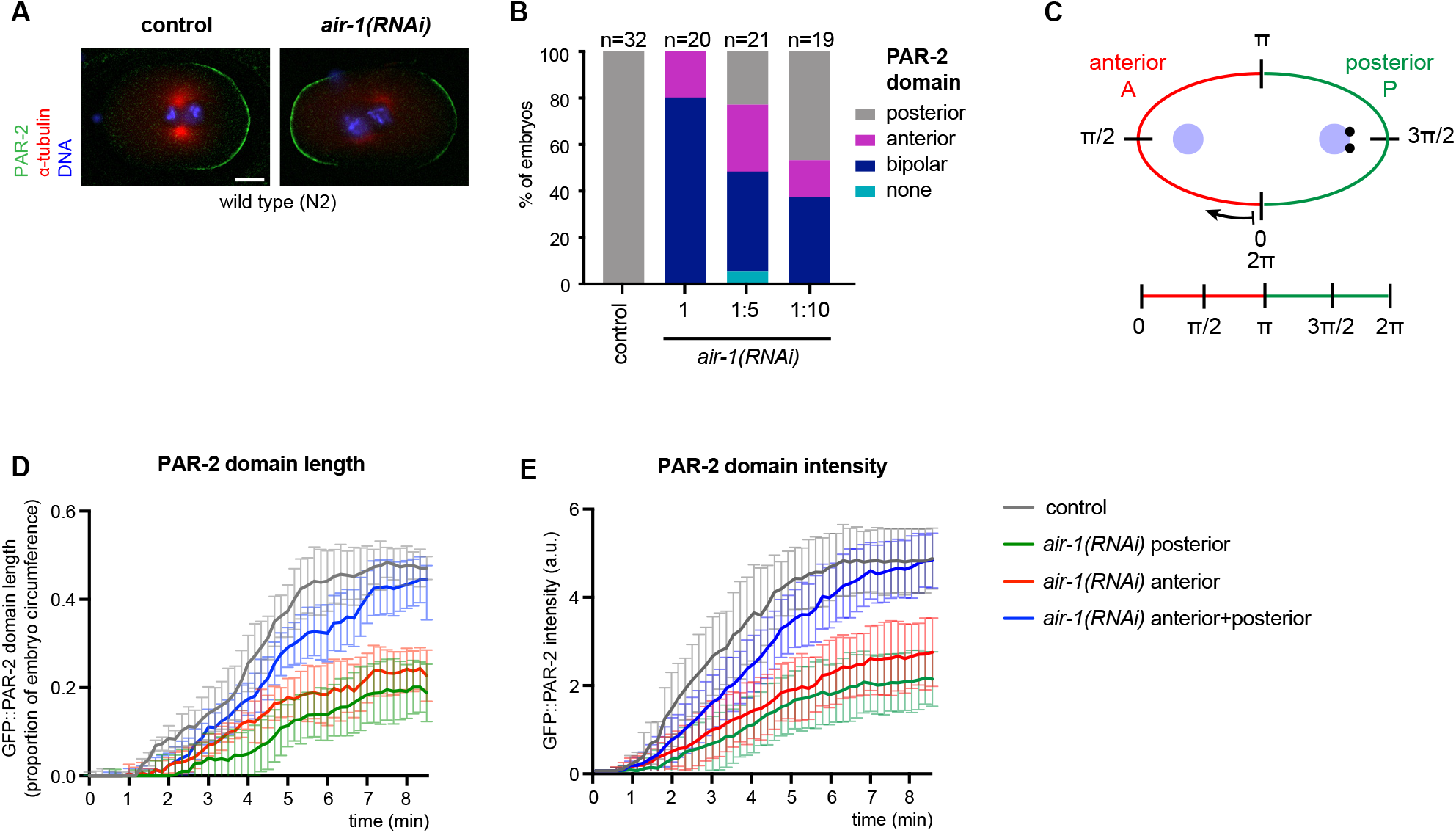
Aspects of polarity in AIR-1-depleted embryos. (A) Control and *air-1*(*RNAi*) embryos at pronuclear meeting, fixed and stained with antibodies against α-tubulin (red), PAR-2 (green); DNA is shown in blue. Here and in all other panels scale bar: 10μM. (B) Quantification of PAR-2 distributions in control and *air-1*(*RNAi*) embryos. *air-1(RNAi)* was diluted 1:5 or 1:10 with bacteria expressing an empty vector (L4440), as indicated. (C) Scheme illustrating how the embryo’s circumference was straightened for Fig. 1E,F; Fig.4F,G and Fig. S4A,B (D, E) Length (B) and intensity (C) of GFP::PAR-2 domains over time in control and *air-1(RNAi)* embryos.

**Fig. S2:**
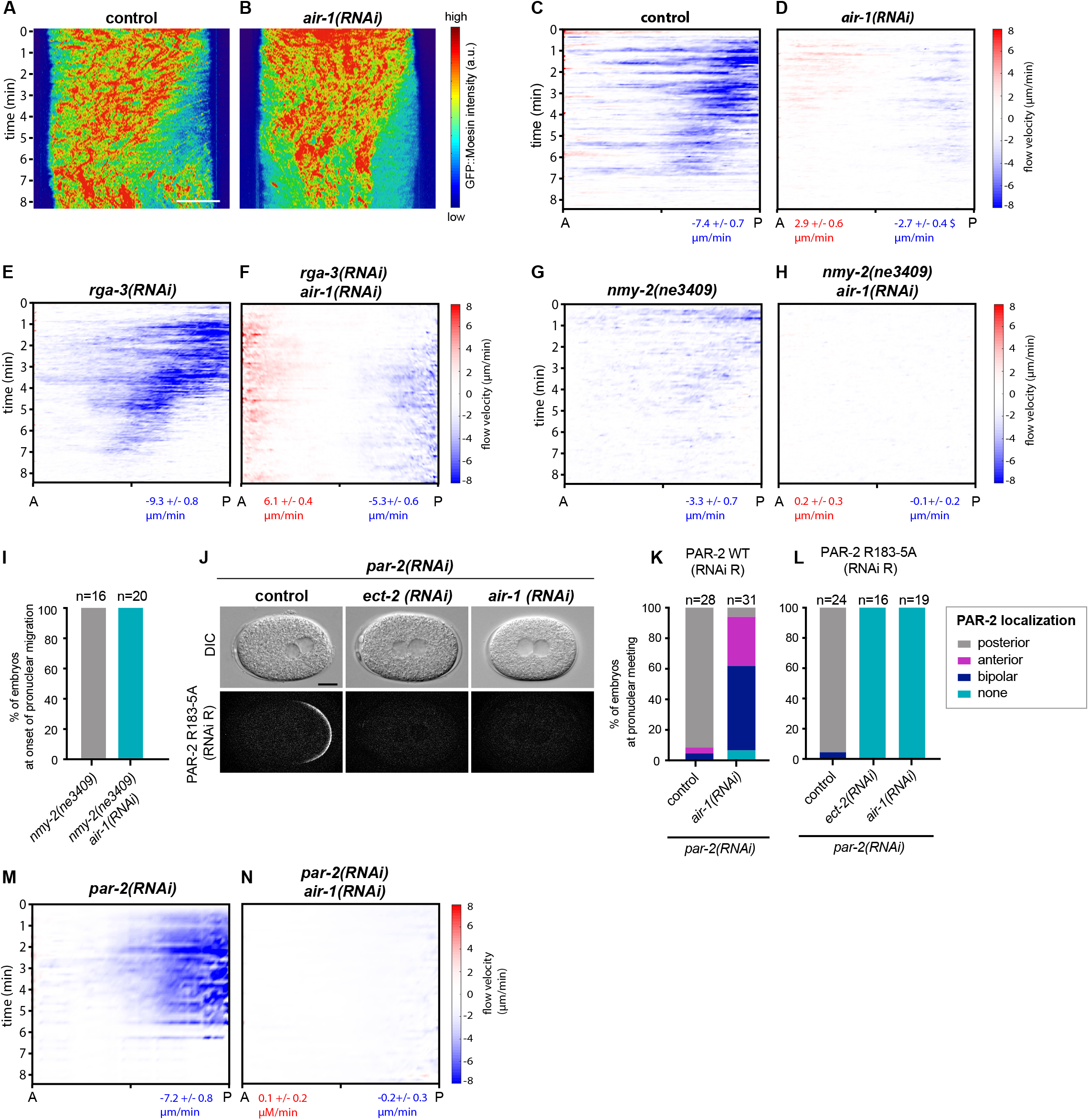
Cortical flows in AIR-1 depleted embryos are PAR-2-dependent. (A,B) Kymographs of control and *air-1*(*RNAi*) embryos expressing GFP::Moesin (show wiht Physics look up table). (C-H) Kymographs of cortical flow velocities quantified by particle imaging velocimetry (PIV) of GFP::Moesin expressing embryos of indicated genotypes. Values indicate average peak velocities at the anterior (red) and posterior (blue). N=5 for all conditions, except for *rga-3(RNAi),* where N=4. (I) Quantification of GFP::PAR-2 distributions in *nmy-2(ne3409)* and *nmy-2(ne3409)+air-1(RNAi)* embryos at the onset of pronuclear migration. (J) Embryos expressing GFP::PAR-2 R163A depleted of endogenous *par-2* either alone or together with *ect-2(RNAi)* or *air-1(RNAi),* as indicated. (K,L) Corresponding quantification of GFP::PAR-2 distributions. (M,N) Kymographs of cortical flow velocities quantified by PIV of embryos shown in M. Values indicate average peak velocities at the anterior (red) and posterior (blue) N=5 for both conditions.

**Fig. S3:**
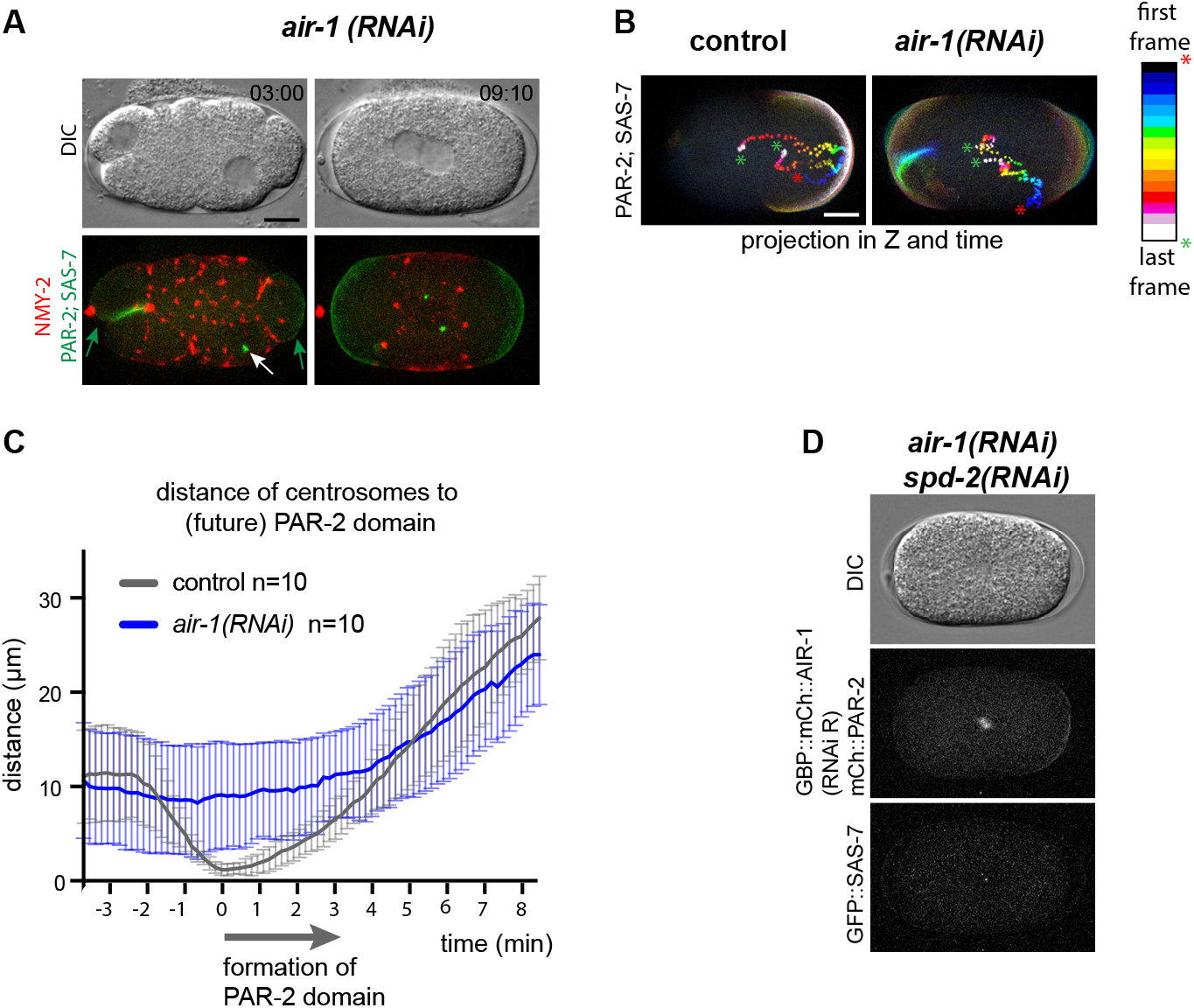
Impaired centrosome positioning in AIR-1 depleted embryos. (A) *air-1*(*RNAi*) embryo at the onset of pronuclear migration expressing GFP::PAR-2 (green), GFP::SAS-7 (green) and RFP::NMY-2 (red). White arrow: centrosomes, green arrows: PAR-2 domains. Note that the posterior PAR-2 domain develops initially at a distance from centrosomes. (B) Projection over time of embryos shown in A. (C) Distance of centrosomes to the PAR-2 initiation site in control and *air-1*(*RNAi*) embryos. (D) *air-1(RNAi) spd-2(RNAi)* embryo at spindle formation expressing GBP::mCh::AIR-1 and mCh::PAR-2. Hermaphrodites were mated with males expressing GFP::SAS-7. Note defective spindle assembly.

**Fig. S4:**
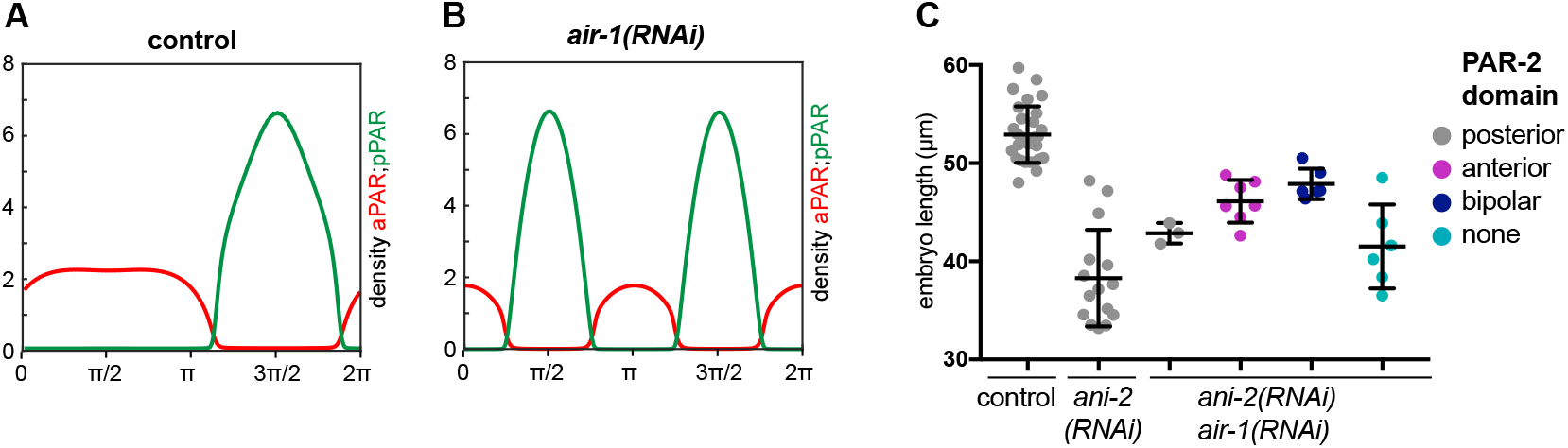
Physical model captures the AIR-1 depletion phenotype. (A,B) Simulation of cortical distributions of anterior (red) and posterior (green) PAR proteins in control (A) and *air-1*(*RNAi*) (C) embryos. (C) Dot plot of size distribution and GFP::PAR-2 domain localization in embryos of indicated genotypes.

## References

1. L. Rose et al. WormBook: the online review of C. elegans biology, 1 (Dec 30, 2014).

2. E. Munro et al. Cold Spring Harbor perspectives in biology 1, a003400 (Oct, 2009).

3. F. Motegi et al. Nature cell biology 8, 978 (Sep, 2006).

4. S. Zonies et al. Development 137, 1669 (May, 2010).

5. F. Motegi et al. Nature cell biology 13, 1361 (Oct 09, 2011).

6. C. R. Cowan et al. Nature 431, 92 (Sep 02, 2004).

7. S. Kotak et al. Journal of cell science 129, 3015 (Aug 01, 2016).

8. J. M. Schumacher et al. Development 125, 4391 (Nov, 1998).

9. E. Hannak et al. The Journal of cell biology 155, 1109 (Dec 24, 2001).

10. M. Toya et al. Nature cell biology 13, 708 (Jun, 2011).

11. N. Portier et al. Developmental cell 12, 515 (Apr, 2007).

12. S. Schonegg et al. Proceedings of the National Academy of Sciences of the United States of America 104, 14976 (Sep 18, 2007).

13. J. Liu et al. Developmental biology 339, 366 (Mar 15, 2010).

14. D. R. Hamill et al. Developmental cell 3, 673 (Nov, 2002).

15. K. F. O’Connell et al. Developmental biology 222, 55 (Jun 01, 2000).

16. A. Bezler et al. Genetics 186, 1271 (Dec, 2010).

17. B. Goldstein et al. Development 122, 1467 (May, 1996).

18. N. W. Goehring et al. The Journal of cell biology 193, 583 (May 02, 2011).

19. A. S. Maddox et al. Development 132, 2837 (Jun, 2005).

20. M. R. Wallenfang et al. Nature 408, 89 (Nov 2, 2000).

21. M. Yan et al. Medicinal research reviews 36, 1036 (Nov, 2016).

## Supplementary References

22. V. Praitis et al. Genetics 157, 1217 (Mar, 2001).

23. D. J. Dickinson et al. Genetics 200, 1035 (Aug, 2015).

24. R. S. Kamath et al. Genome biology 2, RESEARCH0002 (2001).

25. R. S. Kamath et al. Methods 30, 313 (Aug, 2003).

26. J. F. Rual et al. Genome research 14, 2162 (Oct, 2004).

27. S. Pichler et al. Development 127, 2063 (May, 2000).

28. F. Tostevin et al. Biophysical journal 95, 4512 (Nov 15, 2008).

29. S. Seirin Lee et al. Journal of Theoretical Biology 382, 1 (Oct 7, 2015)

30. J. F. Joanny et al. The European physical journal. E, Soft matter 36, 52 (May, 2013).

31. M. Mittasch et al. Nature cell biology 20, 344 (Mar, 2018).

32. M. Mayer et al. Nature 467, 617 (Sep 30, 2010).

